# Dental calculus formation is linked to diet and phylogeny in mammals

**DOI:** 10.1101/2025.08.08.669340

**Authors:** John L. Richards, Markella Moraitou, Ella Nates, Konstantina Saliari, Emmanuel Gilissen, Zena Timmons, Andrew C. Kitchener, Olivier S. G. Pauwels, Richard Sabin, Phaedra Kokkini, Roberto Portela Miguez, Katerina Guschanski

## Abstract

1. As a point of entry, the oral cavity serves as a potential first area of colonization of the host with environmental bacteria. As such, the oral microbial community not only affects a wide range of internal host processes, including health and digestion but may also reflect the external environment and ecology of the host.
2. Dental calculus, a mineralized form of dental plaque, preserves the DNA of the host microbiome through time, serving as a rich source of historical microbiota. Despite its potential to provide temporal information on the ecological and evolutionary processes of microbiomes, dental calculus has rarely been explored outside of humans and non-human primates. Hence, it remains unclear how ubiquitous it is across mammals.
3. Using natural history museum collections, we surveyed > 1600 specimens belonging to 142 species, representative of every mammalian order, to investigate the taxonomic distribution of dental calculus, and to identify factors that most strongly contribute to its formation. We found dental calculus to be abundant across mammalian taxa, with 104 of the surveyed species showing calculus deposits.
4. High fiber diets were most strongly associated with calculus presence and abundance, whereas species with high protein and fat diets showed little to no calculus deposits. We also found evidence of phylogenetic signal in calculus formation, pointing to the effects of oral/dental morphology. In addition, captivity strongly affected dental calculus formation in almost all dietary categories. Using this information, we made predictions about the likelihood of finding dental calculus in unsurveyed mammalian species, with the aim of informing future investigations.
5. Our study found that dental calculus is readily available in natural history museum collections, making it an easily accessible source of oral microbiota from wild animals. We highlight the taxonomic diversity of species presenting dental calculus and provide information and suggestions to researchers and museum curators interested in utilizing dental calculus for the study of oral microbiota, past and present.

## 1 Introduction

### 1.1 Dental calculus and historical microbiota

Dental calculus (DC) is mineralized dental plaque biofilm, which is visible as calcified deposits on teeth, and provides a valuable source of microbial biomolecules preserved through time (Warinner et al., 2015). Dental calculus is formed through regular, sequential deposition and calcification of the microbial biofilm, trapping other molecules present in the oral cavity, digestive and respiratory tracts, and the environment, reflecting its unique position as a contact point between host internal systems and the outside. Entrapped molecules include dietary phytoliths (Armitage, 1975; Power, Salazar-García, Straus, et al., 2015; Power, Salazar-García, Wittig, et al., 2015), bacterial, viral, and fungal DNA of the host oral, digestive, and respiratory microbiomes (Brealey et al., 2020; Fellows Yates et al., 2021; Mann et al., 2018), host and dietary DNA and proteins (Hendy et al., 2018; Mackie et al., 2017; Putrino et al., 2024), as well as bioactive inorganic compounds such as heavy metals (Yaprak et al., 2017). This combination of materials makes dental calculus ideal for studying the interplay of host-associated microbial, dietary, and environmental factors.

The long-term preservation of diverse (bio)molecules in dental calculus permits investigation across wide temporal scales. Analysis of microscopic remains in dental calculus enabled reconstruction of past diets, informing archaeological research in humans, non-human primates, and ungulates (Armitage, 1975; Ciochon et al., 1990; Hendy et al., 2018).

Applications of ancient DNA (aDNA) techniques to historical and ancient metagenomic samples, including preserved microbiomes, have enabled much more comprehensive studies into the microbiota of the past and their coevolution with their hosts (Brealey et al., 2021; De La Fuente et al., 2013; Fellows Yates et al., 2021; Moraitou et al., 2022; Rivera-Perez et al., 2015). Dental calculus and paleofeces are the two forms of host-associated microbiomes that can be preserved through time. Dental calculus has many advantages over paleofeces, as it is far more common, less affected by contamination and decomposition, and can more readily be associated with a host individual (Adler et al., 2013; Warinner et al., 2014). Further, in contrast to paleofeces, the preserved oral microbiome of dental calculus reflects an extended period in the life of the host, likely from months to years, and serves as a representation of historical environments and their effects on host microbiota, as the oral cavity serves as a point of entry for environmental microbes (Brealey et al., 2021; Fellows Yates et al., 2021; Moraitou et al., 2022; Shaiber et al., 2020).

### 1.2 Dental calculus formation

The diversity of information captured and preserved through time in dental calculus makes it ideal for addressing a large range of research questions in ecology and evolution and to investigate environmental changes affecting the host species. Despite this great potential, little is known about which species produce dental calculus, and there is a lack of basic knowledge of the factors of host ecology and biology that contribute to its formation. Dental calculus has been studied in several mammal species outside of humans (Armitage, 1975; Brealey et al., 2020, 2021; Moraitou et al., 2022; Ozga & Ottoni, 2023; Power, Salazar-García, Straus, et al., 2015; Power, Salazar-García, Wittig, et al., 2015), but has primarily been considered as a health concern, with the literature on its formation and development being dominated by veterinary studies in pets and livestock (Clarke & Cameron, 1998; Hernández-Castañeda et al., 2015; Verstraete, 2003).

In the development of DC, the formation of a biofilm in the oral cavity is first dependent on the colonization of the tooth surface by microorganisms sustained by nutrients present in the oral fluid (Lieverse, 1999). The presence of salivary amylase is thought to be important in the development of this initial biofilm layer, as it breaks down starches into simple sugars that form a film adhering to the surface of the tooth. Further fermentation of salivary sugars into acids breaks down the tooth enamel, providing more surface area for biofilm adhesion (Radini et al., 2017). Mineralization occurs through the deposition of calcium phosphate crystals in the biofilm cellular matrix, and the crystal structures strengthen and mature through time, serving, once mineralized, as a surface onto which a new biofilm layer can subsequently adhere and eventually mineralize (Akcalı & Lang, 2018).

A range of factors have been proposed to contribute to dental calculus formation, including nutrients, pH, water content, and salivary flow (Jin & Yip, 2002; Lieverse, 1999). Conflicting reports have attributed heavy calculus formation to high fat, high protein, and high carbohydrate diets, while also highlighting the effects of salivary calcium, phosphate, and pH (see Lieverse, 1999). Other factors have the potential to impact dental calculus formation, including tooth morphology, shape of the oral cavity, host behavior, and location of salivary glands (Radini et al., 2017). Many aspects of morphology and feeding behavior that may impact dental calculus formation are strongly coupled with diet but also to host taxonomy, e.g., similarities in tooth shape may reflect similarities in diet but also be constrained by evolutionary relationships between hosts (Evans & Pineda-Munoz, 2018).

### 1.3 Dental calculus in natural history collections

Natural history museum collections are ideally suited for the study of dental calculus, as skulls are among the most frequently collected specimens. Dental calculus deposits develop on the surface of the tooth and hence their sampling does not damage the underlying tooth morphology. However, like any destructive sampling, once removed and processed, calculus cannot be recovered. Therefore, both researchers and curators would benefit from a better understanding of which species are likely to reliably form dental calculus, so that specific research questions and efficient sampling strategies can be developed for the use of this precious material.

To support researchers in planning their studies and museum curators in accommodating and anticipating sampling requests, we aimed to i) assess which mammal species develop dental calculus, and to what extent and ii) uncover underlying principles of dental calculus formation in mammals by considering ecological and phylogenetic aspects of mammalian biology. Lastly, using collected data, we make predictions about dental calculus presence based on host taxonomy and dietary composition, looking towards future developments in the use of dental calculus to study historical microbiota. The potential of dental calculus in reconstructing (past) oral microbiota is immense, and in exposing the huge range of as-yet unexplored sources of dental calculus in a large diversity of species, we hope to highlight the taxonomic and ecological diversity of mammalian host taxa that may be studied.

## 2 Methods

### 2.1 Dataset and Calculus Survey

To assess the prevalence of dental calculus formation across mammals, we surveyed at least one species in each mammalian order, for a total of 1629 specimens belonging to 142 species in 129 genera across 72 families (Table S1). Surveyed specimens were housed at the National Museums Scotland, Edinburgh, UK (NMS); the Natural History Museum London, UK (NHMUK); the Naturhistorisches Museum Wien, Austria (NHM Vienna); the Royal Belgian Institute of Natural Sciences, Brussels (RBINS), and the Royal Museum for Central Africa, Tervuren, Belgium (RMCA). Species names were recorded as they appeared on specimen labels (Table S1 “Recorded Species”). Captivity status was also taken from the labels; for the purposes of our analysis, we treated any indication of time spent in captivity as “captive”. Teeth were scored and classified according to the presence and abundance/amount of calculus (Fig. 1), with 0 = no calculus; 1 = minimal amount of calculus present across few teeth (e.g., 1-2 isolated plaque deposits); 2 = several large plaque deposits across multiple teeth or most teeth with similar low amount of calculus; 3 = total encrustation of several teeth or large plaques on most teeth.

**Figure 1:**
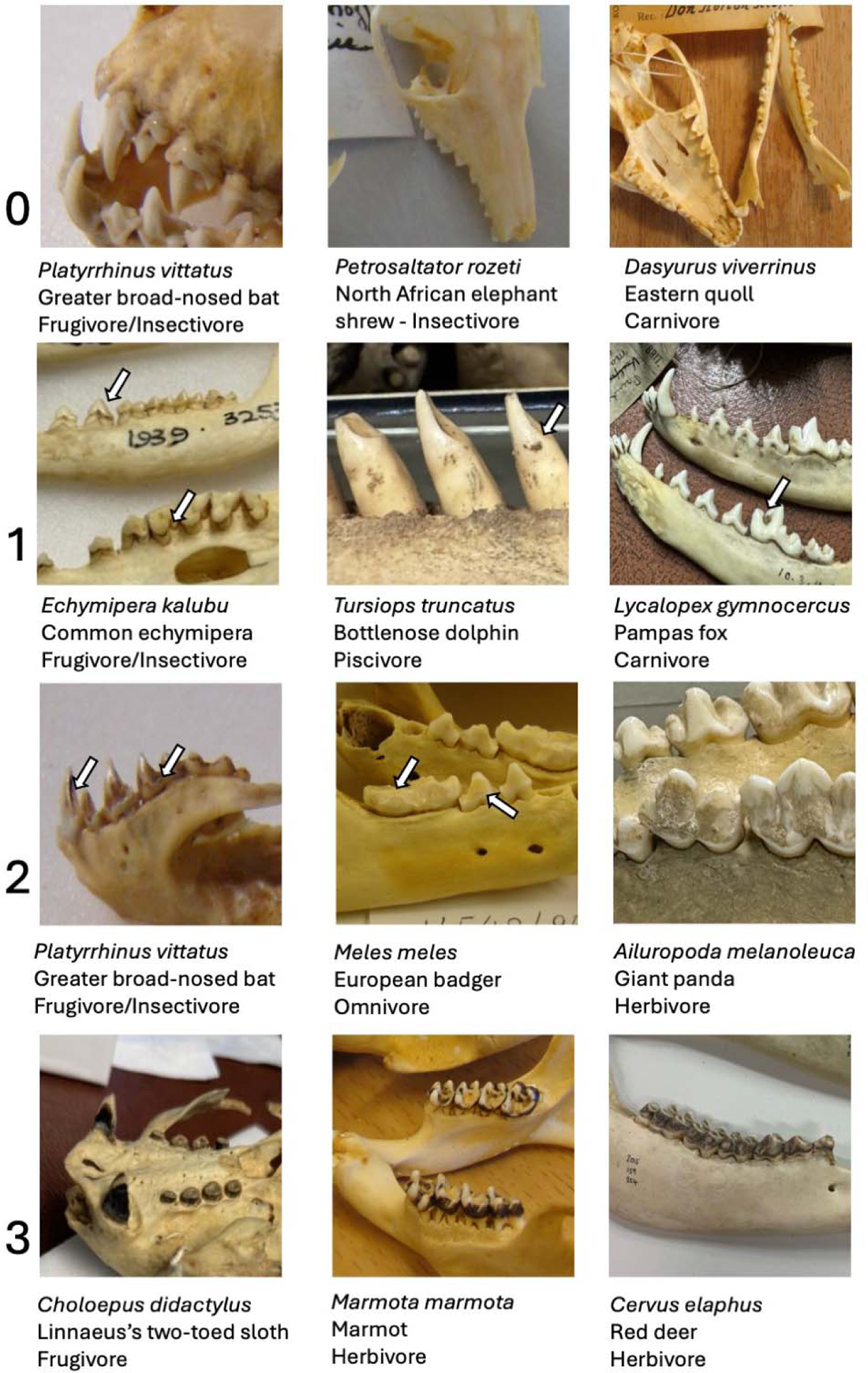
Examples of specimen dental calculus (DC) scores. DC scores given (on the left, 0-3) for 11 species of mammal. Common names and dietary classification according to Lintulaakso et al., 2023 provided for each specimen. Arrows indicate calculus deposits. Two specimens of *P. vittatus* with different scores are shown.

Teeth were scored separately for upper and lower jaws (maxillae and mandible), lingual and buccal tooth surface, and tooth type (incisor, canine, and [pre]molar). We aimed to survey 10 specimens per species when possible, though due to collection limits and missing teeth, not all species were represented by 10 fully intact specimens (Table S1). When possible, we aimed to survey geographically and temporally diverse specimens to avoid a potential batch effect resulting from museum preparation practices or environment. Only teeth set in the jaw were surveyed to increase the confidence in tooth type assignment and specimen identity. Teeth not set in the jaw, even if showing a clear tooth type morphology and association with a specimen (e.g., in the same box), were disregarded. Cetaceans have a homodont tooth morphology and these teeth were all classed as molars. Statistical and comparative analyses were carried out only on species for which four or more specimens were surveyed, for a total of 118 species, 110 genera, and 64 families, short only one order, Hyracoidea. However, the majority of species had 10 specimens surveyed (n=72). The results of the survey, including species with fewer than four individuals, are presented in Supplementary Materials Table S1.

We summed and averaged scores across the specimens surveyed per species to get a mean score per tooth type/side/jaw, referred to as DC score (Table S2). We used an ANOVA to test for the effects of tooth type, side, and jaw on DC scores. Because only the tooth type was significant (see below), we estimated a mean score per tooth type for each species by averaging upper and lower lingual and buccal scores for the given tooth type across all individuals. We used molar DC scores for all subsequent analyses, unless specified otherwise, as molars were the most consistently present tooth type in the study specimens and thus the datasets had the least amount of missing data (see below).

To test for differences in DC score related to diet, we assigned surveyed species into diet groups following the “calculated cluster mean diet” classification from Lintulaakso et al., 2023. We used an ANOVA to test for differences in molar DC score between the four diet groups: animalivore (n=48), herbivore (n=31), frugivore (n=18), and omnivore (n=21).

To rule out the possibility that different preservation approaches and specimen preparation techniques could have affected dental calculus prevalence and abundance in different museums, we selected five species with at least four surveyed specimens surveyed in two different museums and tested for the consistency in DC scores between the museums for the same species using a nested ANOVA. We included species with different diets (animalivorous *Panthera leo* and *Puma concolor,* frugivorous *Pteropus vampyrus,* and herbivorous *Capreolus capreolus* and *Galeopterus variegatus*) to account for dietary variation.

To assess the effects of captivity on DC formation, 71 surveyed species contained either captive or both captive and wild individuals, with a total 466 specimens that had been held in captivity. We used nested ANOVAs to test for differences in DC score between captive (primarily from zoos) and wild individuals across diet groups.

### 2.2 Phylogenetic analyses

We tested for phylogenetic signal in dental calculus formation with two approaches implemented in the R packages picante and phylosignal (Cowan et al., 2020; Keck et al., 2016). Since one of the approaches relies on binary data, we converted DC scores into presence/absence records by collapsing mean DC scores per tooth type ≤ 0.1 to 0 (absence of DC) and mean scores >0.1 to 1 (presence of DC). The cut-off corresponds to a single specimen out of 10 having a maximum DC score for a specific tooth type (e.g., molar) on each surface (buccal and lingual) and jaw (upper and lower), or all specimens consistently showing a DC score of 1 for a specific tooth type on each side and jaw. We downloaded the MCC mammalian consensus tree constructed with DNA-only records from vertlife.org (Upham et al., 2019). Five species we surveyed were not present in this tree. For three of them (*Anomalurus derbianus, Tupaia montana,* and *Oryzorictes tetradactylus*) we used a congeneric species as representative (*A. beecrofti, T. glis,* and *O. hova*). One species (*Elephantulus fuscipes*) was excluded from analysis as two congeneric species were already present, and one monotypic species (*Lavia frons*) was represented by its closest relative (*Megaderma spasma*).

The first phylogenetic approach utilized net-relatedness index (NRI) and nearest taxon index (NTI). Both indices are measures of phylogenetic clustering or overdispersion and as such can be used as a measure of phylogenetic signal. The NRI is a measure of the difference in mean pairwise distance (MPD) between all species pairs with the trait present compared to random reshuffling, and so indicates whether traits are found more clustered than expected. The NTI is similar, but uses the mean nearest taxon distance (MNTD) between all species with the trait; the NTI shows if any two species with the trait are closer to each other than would be expected if the trait was randomly distributed.

Second, we used Pagel’s lambda for the continuous trait values of mean DC score per tooth type (Gittleman & Kot, 1990; Pagel, 1999). Pagel’s lambda is a measure of the transformation of a phylogeny that fits the trait data to a model of Brownian motion. A lambda value of 1 means the trait is distributed as expected under Brownian motion, and 0 is random (equivalent to a star phylogeny).

### 2.3 Diet analysis

We assigned surveyed species into diet groups following the “calculated cluster mean diet” classification from Lintulaakso et al., 2023. To test for differences in molar DC score related to diet, we used an ANOVA for four diet groups: animalivore (n=48), herbivore (n=31), frugivore (n=18), and omnivore (n=21). The same publication was also used to obtain dietary nutritional composition for the surveyed mammalian species (Lintulaakso et al., 2023). This database lists diet nutrient composition per species divided into five components: crude fiber, inorganic residue (ash), crude fat (ether extract), protein, and sugars and starches (nitrogen-free extract). We followed the tripartite dietary scheme in Lintulaakso et al., (2023), combining crude fiber and ash, crude protein and fats, and sugars and starches, roughly corresponding to herbivores, animalivore, and frugivores, respectively.

To disentangle the effects of diet and phylogeny on DC formation, we used a multilevel linear mixed effects model corrected for phylogenetic relatedness implemented in the R package Metafor (Viechtbauer, 2010). We fitted diet nutritional composition as the explanatory variable and molar DC score as the response variable. As the diet data was compositional in nature (all nutrient components summed to 1), log-ratios were used. Each of the three log-ratios was used as a fixed predictor variable: Protein & Fats:Fiber; Sugars & Starches:Fiber; Protein & Fats:Sugars & Starches. To account for the effects of phylogeny, we included taxonomic species as a random effect and phylogeny as a variance-covariance matrix derived from the tree used above. To account for measurement errors, the squared standard error for each species’ molar DC score was included as *V*.

### 2.4 Predictions of DC scores across the mammalian tree of life

The model constructed above was used to make predictions of molar DC scores across ∼4,500 mammal species present in the diet composition database but not surveyed in this study. The species-specific predicted DC scores were averaged by mammalian family to provide an estimate of the likelihood of dental calculus presence.

## 3 Results

### 3.1 Dental calculus is present in majority of mammals

We identified DC as the presence of a visible deposit on the teeth of surveyed specimens that was clearly distinct from any tooth morphological features (Fig. 1). The DC deposits varied in form, size, and color, ranging in appearance from chunky white plaques to fine dark film. They were often confined to what was likely a supragingival position on the living individuals (Fig. 1, examples for scores 2 and 3), supporting their identification as DC. To further verify that these varied deposits were indeed DC, an associated study sequenced samples from 403 individuals in 30 mammalian species and confirmed the presence of oral microbial taxa in all samples (Moraitou et al., 2025). We can thus be confident that here we report on dental calculus, the calcified oral microbiome, from museum specimens.

We found no significant differences in DC scores for jaw (upper versus lower, t(1158) = −0.32, p = 0.748, Fig. S1A) or side (lingual versus buccal, t(1156) = −0.60, p = 0.551, Fig. S1B) within the entire dataset. However, the presence of calculus varied by tooth type.

Specifically, molars most frequently and consistently showed DC deposits, so that if a species produced DC, it was present on the molars but was often absent on other teeth (Fig. 2). DC scores differed significantly across tooth types (ANOVA, F(2, 1157) = 111.76, p < .001), with molars consistently showing higher DC scores (Fig. 3A). Post hoc Tukey’s HSD tests showed that molars had significantly higher scores than both canines and incisors, but there was no significant difference between incisors and canines (Table S3A).

**Figure 2:**
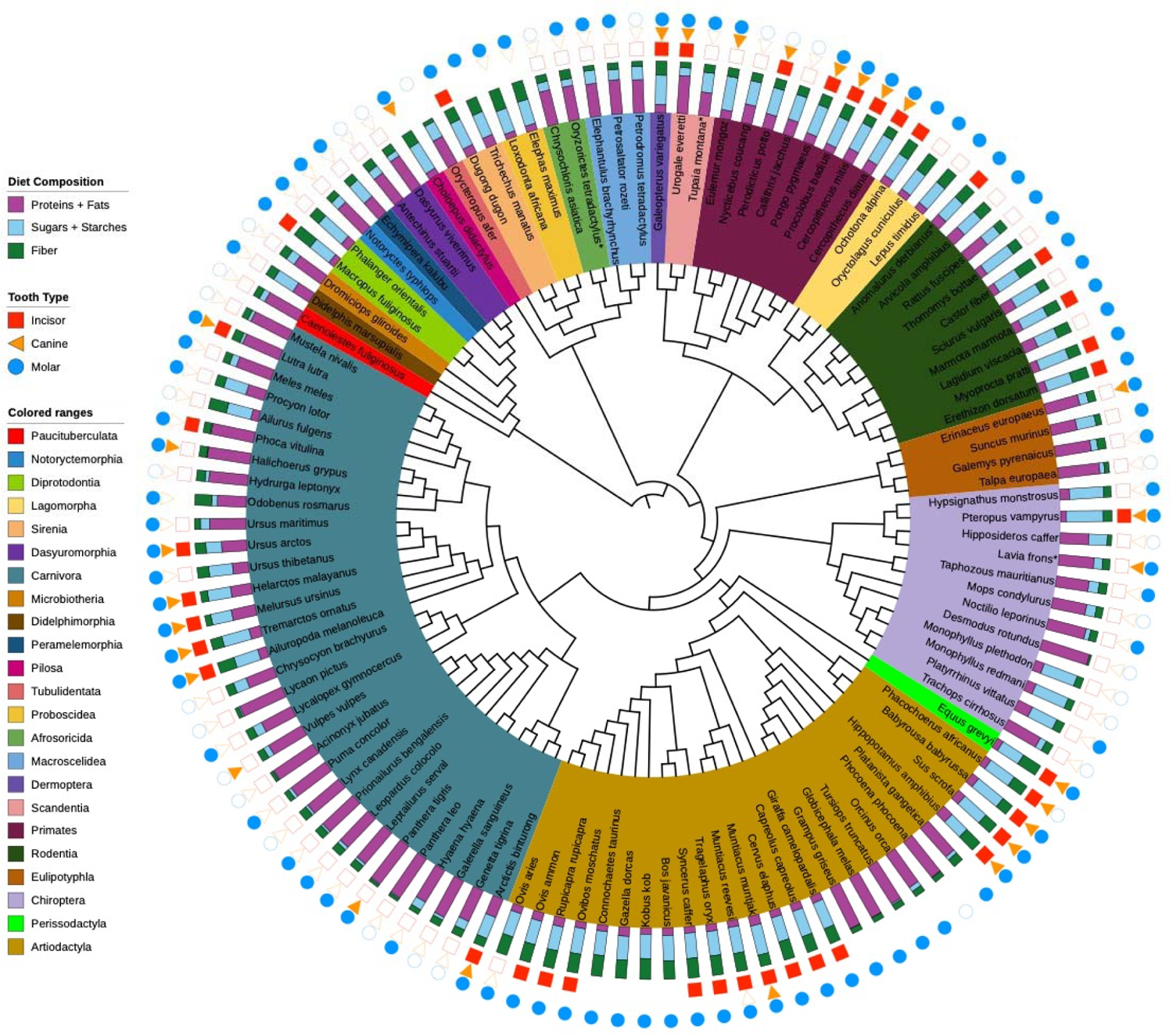
Dental calculus presence across the mammalian phylogeny and tooth types, showing species for which at least four wild specimens were available (n=118). Mean scores per tooth type were binarized, with a score >0.1 considered as DC presence and shown with a filled shape along the three outer rings. Unfilled shapes indicate a mean score ≤0.1, which was considered as absence of DC. No shape indicates that the tooth type was absent in the specimens surveyed or that the species lacks that tooth type altogether. All cetacean teeth were scored as molars. This tree was downloaded from vertlife.org (Upham et al., 2019) using the MCC consensus tree of DNA-only records. Asterisks indicate surveyed species not present in the downloaded tree. For *Anomalurus derbianus, Tupaia montana,* and *Oryzorictes tetradactylus* we used congeneric species as representative: *A. beecrofti, T. glis, O. hova*.

**Figure 3:**
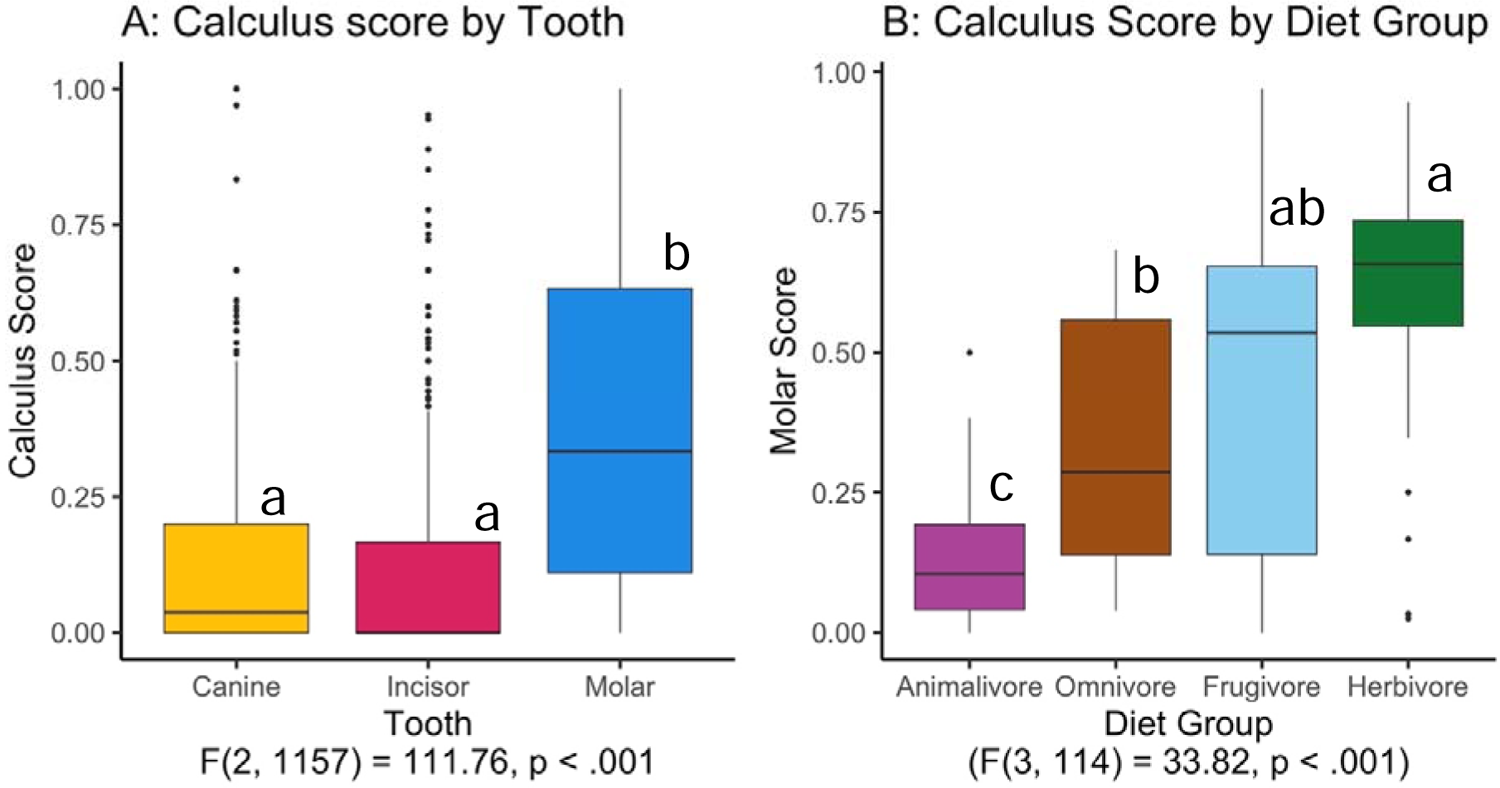
Comparisons of calculus scores by tooth type (A) and molar calculus scores by diet group. **(B)**. Scores for wild specimens only. Results from one-way ANOVAs are shown below X axis label. Letters above the boxplots indicate results from post hoc Tukey’s HSD tests. In A: only molars differ significantly from canines and incisors. In B: frugivores, herbivores, and omnivores all show significantly higher scores compared to animalivores. No significant difference was found between herbivores and frugivores, or frugivores and omnivores. Omnivores have significantly lower scores than herbivores. Values for post hoc tests can be found in Table S3B.

Additionally, molars were the most reliably present tooth type on the surveyed specimens, with many specimens missing incisors or canines but rarely missing molars (Table S1). Only three species did not follow this general pattern (*Perodicticus potto, Vulpes vulpes,* and *Phacochoerus africanus*), as they showed DC on incisors and/or canines, but not on molars (Fig. 2, Table S1). Owing to the consistency of DC on molars, we aggregated DC scores for jaw and side, as described in the Methods, and focused all subsequent analyses on molar DC scores.

Of the 118 surveyed species with four or more specimens, 104 presented dental calculus, most commonly on the molars (Fig. 2, Fig. S2 for survey results including all 142 species, which were qualitatively similar). Calculus abundance varied within and across species (Table S1). Some species presented dental calculus in large accretions, regularly developing on the same tooth types. Other species consistently showed sparse calculus presence (Table S1). Few species showed absolutely no calculus (Fig. 2, Fig. S2). To evaluate if the obtained DC scores could be due to differences in specimen preservation and treatment among different natural history collections, we identified five species for which at least four specimens were surveyed in two different museums each. DC scores did not differ for four out of five species, whereas we observed a significant difference in DC scores for *Puma concolor* (Fig. S1D). However, this difference may have an ecological explanation (see Discussion). In general, this result suggests that the observed DC scores are a biological feature of the surveyed species and not primarily influenced by preservation, treatment, and storage conditions in the museums.

### 3.1 Production of dental calculus is phylogenetically constrained

Next, we aimed to investigate any underlying phylogenetic structure to dental calculus formation. First, we used presence-absence values for DC. NRI and NTI measure the effect size of MPD and MNTD, representing evolutionary relationships at a deep and shallow (tip to tip) level, respectively. NRI values of 1.96 and −1.96 have been used as thresholds beyond which traits are considered phylogenetically clustered or overdispersed, respectively (Coppi et al., 2019). For all tooth types, our results show clear evidence of clustering, with NRI values all statistically significant and >2 (Table 1). No NTI values were significant, indicating that the distance from any species with DC to its nearest neighbor with DC was not different from what would be found if the trait were distributed randomly.

**Table 1:** Mean Pairwise Distance (MPD), Mean Nearest Taxon Distance (MNTD), Net Relatedness Index (NRI), Nearest Taxon Distance (NTI) values for all three tooth types. Numbers in parentheses indicate the number of species presenting dental calculus for each tooth type under the binarized scoring scheme out of the number of species with that tooth type. Column Observed shows values calculated from our data, Random Mean and Random SD show the mean and standard deviation from the null model of 999 random tip reshufflings. NRI and NTI measure phylogenetic clustering or overdispersion, equal to the standardized effect size multiplied by −1. Asterisks indicate *p*<0.05.

| TOOTH |  | OBSERVED | RANDOM<br>MEAN | RANDOM SD | NRI<br>NTI |
| --- | --- | --- | --- | --- | --- |
| **MOLAR**<br>**(88/118)** | MPD | 165.45 | 176.87 | 5.61 | 2.03* |
|  | MNTD | 48.9 | 48.6 | 3.2 | 0.11 |
| **CANINE**<br>**(28/82)** | MPD | 141.23 | 177.49 | 15.13 | 2.39* |
|  | MNTD | 61.00 | 74.04 | 9.60 | 1.35 |
| **INCISOR**<br>**(39/108)** | MPD | 145.21 | 176.87 | 10.99 | 2.88* |
|  | MNTD | 56.13 | 63.06 | 6.60 | 1.05 |

Second, we used Pagel’s lambda to test for the presence of phylogenetic signal in molar DC score. Pagel’s lambda ranges from 0 to 1, with 1 representing an underlying phylogenetic structure following a model of evolution under Brownian motion. We recovered a value of 0.83 (*p*<0.05) using species molar DC scores, confirming the presence of phylogenetic signal.

### 3.2 Diet and evolutionary relationships impact dental calculus formation

DC formation is expected to be influenced by diet (Jin & Yip, 2002; Lieverse, 1999; Radini et al., 2017). We therefore considered if mammals belonging to different dietary groups differ in the build-up of DC. Indeed, we found a significant difference in DC scores among mammals by dietary groups, with herbivores showing higher DC scores than animalivores and omnivores (F(3, 114) = 33.82, p < .001, Fig. 3B). Frugivores, herbivores, and omnivores had significantly higher scores than animalivores in post hoc Tukey’s HSD tests; the difference between herbivores and frugivores was not significant (*p* = 0.058) and similarly, omnivores did not significantly differ from frugivores (*p* = 0.529), but had lower scores than herbivores (Table S3B).

As captivity affects diet (among many other aspects), we also tested for differences in DC scores between wild and captive individuals in our dataset. Indeed, captivity had a significant effect on the DC scores (nested ANOVA, F(6, 856) = 57.03, *p* < .001, Fig. S1C). Molar scores for captive individuals were more similar to each other across diet groups than for wild individuals (Fig. S1C), suggesting a homogenizing effect of captivity. Specifically, average molar DC scores for species with animalivorous diets were higher in captivity than in the wild, whereas average molar DC scores for species with herbivorous and frugivorous diets were lower.

We next examined which nutritional components affect DC formation. To this end, we used the tripartite dietary nutrition structure in Lintulaakso et al. (2023), reflecting the main diet groups: proteins and fats for animalivory, sugars and starches for frugivory, and fibers for herbivory. DC scores were highest for species with low protein and fat content in their diets (Fig. 4), mirroring the analyses based on dietary group assignments above. However, the presence of outliers – species that did not form DC although their dietary composition is suitable for DC formation and *vice versa* (Fig. 4) – suggested that factors other than dietary nutritional components affect DC build-up.

**Figure 4:**
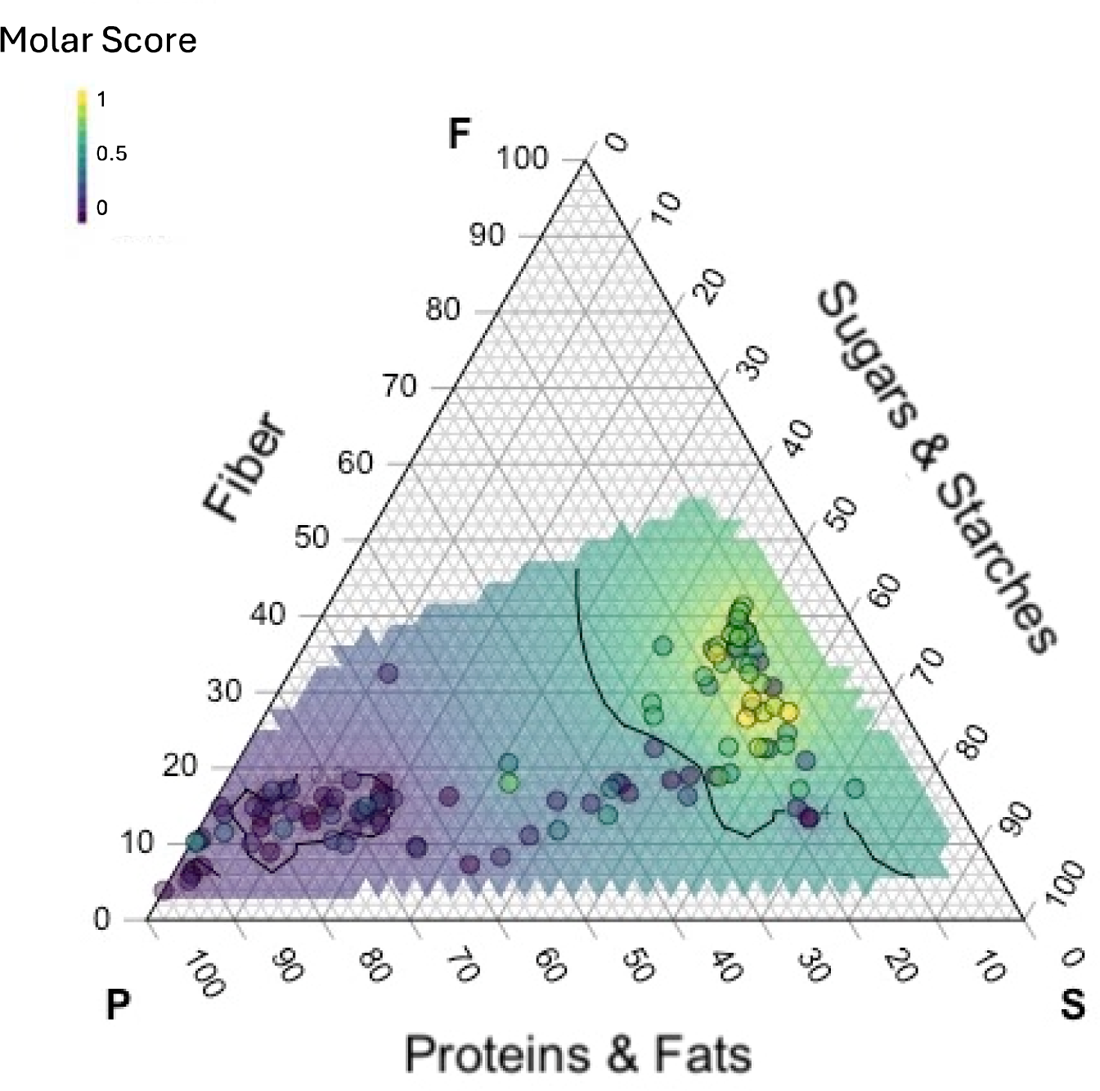
Ternary plot of molar calculus scores and diet nutritional composition. Each dot indicates one species included in this survey. The shaded area covers the full range of diet nutritional composition data taken from Lintulaakso et al. (2023). Letters in the corners indicate the point of maximum value for each nutritional component: F: fibers, S: sugars & starches, P: proteins. The color of the area indicates the projected molar calculus score based on our survey results. Colored dots indicate the scores from surveyed species. Contours delineate areas in diet space with high (right) and low (left) expected DC formation based on diet nutritional composition.

Diet is associated with morphological traits like tooth shape. Indeed tooth shape has been used to infer diet (see Evans & Pineda-Munoz, 2018), and has been found to show some degree of phylogenetic signal (Reuter et al., 2023). Therefore, the phylogenetic signal we recovered for DC formation could directly reflect dietary nutritional composition but also other phylogenetically structured factors, such as oral and digestive tract morphology and physiology. We aimed to separate the factors leading to DC formation resulting from the nutritional components of diet, and those resulting from other biological/ecological variables that may also have an underlying phylogenetic structure. To achieve this, we constructed a multilevel linear mixed-effects model that takes both host phylogeny and dietary nutritional composition into account. The model returned a much stronger effect of protein-to-fiber ratio than protein-to-sugars ratio, with the fibers-to-sugars ratio removed as its effect was redundant and negligible (Table 2). After accounting for the effect of diet on DC formation, the remaining variance was split approximately evenly between taxonomic species and phylogeny, reflecting shallow and deep evolutionary scales, respectively (Table 2).

**Table 2:**
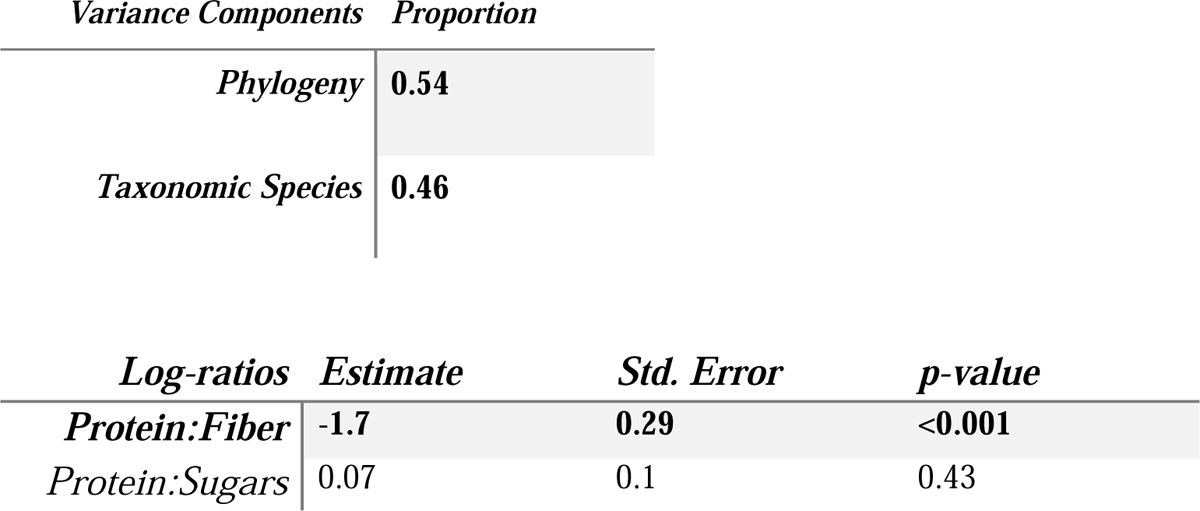
Phylogenetically-informed diet model. Log ratios were used to compare the effects of dietary nutrient proportion. The effect of Fibers:Sugars was negligible and redundant. Variance components Phylogeny and Taxonomic Species were included as random effects of a phylogenetic distance matrix and phylogeny-agnostic taxonomic groups, respectively.

### 3.4 Dental calculus predictions

To facilitate oral microbiome research across the diversity of mammals, we used the insights obtained here to make predictions about the presence and abundance of dental calculus across the mammalian tree of life. Using the above model (Table 2) and dietary information from Lintulaakso et al. (2023), we made predictions about dental calculus scores for ca. 4,500 mammal species that were not included in the survey (Table S4). These predictions were then combined, taking the mean molar DC score value per mammalian family to provide indications for clades that are most likely to present calculus (Fig. 5).

**Figure 5:**
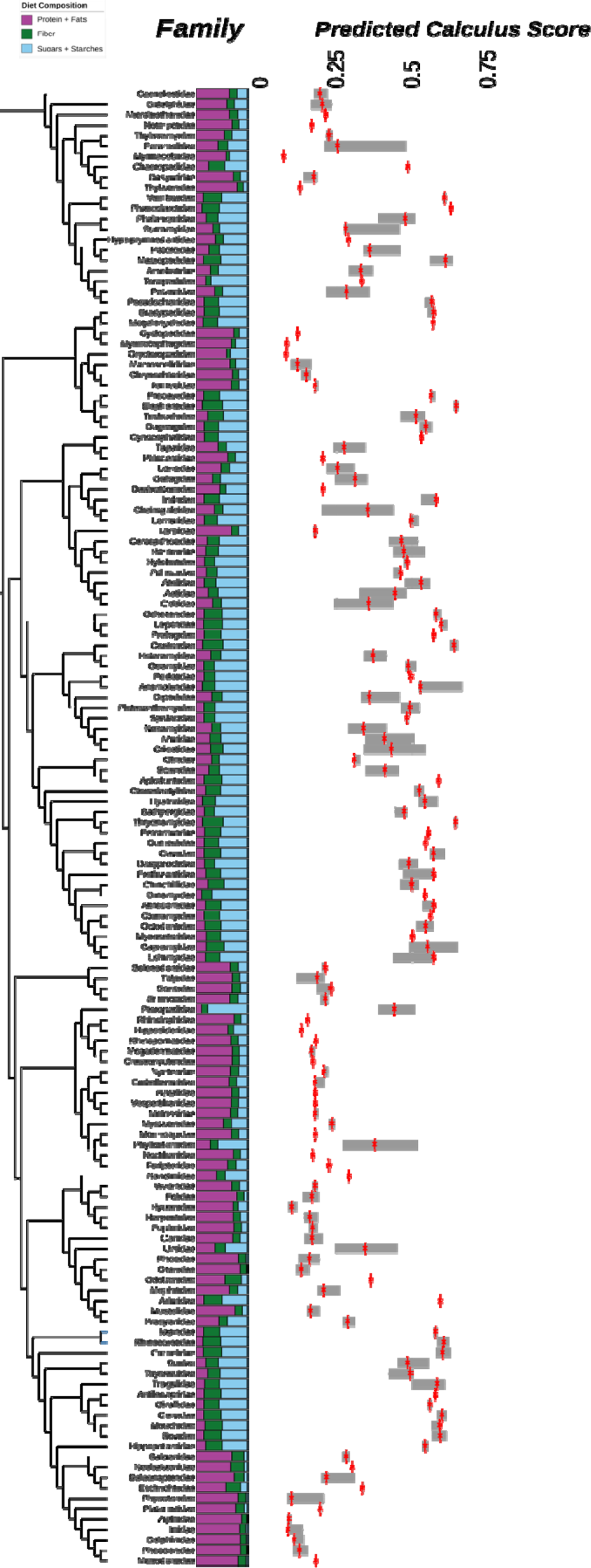
Predicted molar calculus scores by mammalian family. Phylogenetic tree of mammalian families, showing mean diet compositions (bars) and mean predicted calculus score (boxplots) per family. Mean value per family indicated by red bars. This tree was downloaded from vertlife.org (Upham et al., 2019) using the MCC consensus tree of DNA-only records.

## 4 Discussion

### 4.1 Dental calculus survey

Although the use of DC in addressing ecological and evolutionary questions of (historical) oral microbiota and host species is increasing, it has so far been limited to only a few mammalian species, with the focus on primates (Brealey et al., 2021; Fellows Yates et al., 2021; Moraitou et al., 2022; Ozga & Ottoni, 2023; Power, Salazar-García, Wittig, et al., 2015). Our comprehensive survey across over 1,600 wild and captive specimens in five natural history museums demonstrated that DC is present in a large diversity of mammals, in all major mammalian orders, dietary groups, life histories, and habitats. By scoring DC presence and abundance, we revealed some general rules for the development of this calcified oral microbiome in mammals.

We found that DC was most reliably present on molars, even though many species also showed calculus deposits on incisors and canines. Molars are the most abundant tooth type retained in the jaw of dried preserved specimens in natural history collections, enabling reliable association between the individual specimen and tooth. Additionally, molars are used for chewing as opposed to biting or tearing. Therefore, they may experience less abrasion during the mastication process and their position in the mouth may allow for saliva and particulate matter to accumulate and remain in their vicinity for longer than for other tooth types, potentially promoting dental calculus build-up.

Museum preparation practices certainly varied widely from institution to institution and through time, as methods improved and the human ideal of a perfect specimen changed. Unfortunately, records on how individual specimens were prepared are rarely available, making it extremely difficult to evaluate how different practices may have affected retention of dental calculus on the specimen teeth. We avoided specimens that showed obvious evidence of tooth cleaning, e.g., excessive glue affixing teeth, polished bone, varnished tooth surfaces, etc. though of course it is impossible to confidently identify specimens that may have had calculus deliberately removed. Past accidental calculus removal in some specimens is also almost guaranteed, considering decades of handling, moving, and manipulation.

However, for those species for which we had four or more specimens in at least two different museums (species which included representatives of every diet group), we found no significant difference in calculus scores, indicating that differences in specimen treatment and preparation among museums were not impacting the results of our survey. The only exception to this was *Puma concolor* (Fig. S1D). This species has a large geographic range covering the Americas. The majority of samples from this species surveyed at NHMUK (six of seven) were from South America, whereas the majority of samples at NMS (four of six) were from North America, corresponding to two different subspecies living under different ecological conditions (Culver, 2000). Therefore, although differences in specimen treatment and preservation cannot be excluded, there could be a biological explanation for the observed differences in DC scores. Further, in a species with a virtually identical dietary nutritional composition (*Panthera leo*, see diet composition in Table S2), we found no significant differences between museums (Fig. S1D). We fully acknowledge that individual age (rarely recorded, though the majority of individuals for which age information was available were recorded as adults, Table S1), preparation methods, storage conditions and handling routines could result in variation in DC scores and hence our results, particularly about DC absence or low scores, should be treated with caution. Nevertheless, we generally observed good correspondence in DC scores from different collections and highlight the possibility that differences in ecology, as a result of geographic range, or age of the individual could contribute to the reported differences in *Puma*.

### 4.2 Diet and phylogeny explain DC formation

Comparing DC formation between diet groups, we found that herbivores and frugivores consistently developed more calculus than animalivores. This is in line with expectations based on the literature, with carbohydrates and starches frequently reported as contributing to DC formation (Akcalı & Lang, 2018; Lieverse, 1999).

We found that captivity strongly affected DC abundance, with DC scores being more similar among captive individuals belonging to different dietary groups than among wild individuals. We followed the captivity designations from specimen labels. While time in captivity may have varied, we chose to consider any evidence of time in captivity as potentially influencing diet and hence DC formation. Captivity most strongly affected DC formation in species with animalivorous diets, with captive individuals showing higher DC scores and hence more calculus than wild individuals (Fig. S1C). Processed diets, often supplemented with commercial pet food fed to big cats in captivity have been shown to have higher carbohydrate content compared to wild whole-carcass diets (Dzierga, 2014; Kapoor et al., 2016; Kerr et al., 2013; Whitehouse-Tedd et al., 2015), which likely leads to a greater development of DC. On the other hand, herbivores and frugivores both showed decreases in DC formation in captivity, likely due to supplemental crude protein and fat in their feeds (often exceeding 30% in total) leading to reduced calculus formation (Fens & Marcus Clauss, 2024). Mechanical properties of food in captivity also likely have an effect on DC formation. Previous studies in animalivores have shown that softer diets can lead to plaque build-up and disease, as captive diets lack grit that abrade teeth during mastication (Crates et al., 2023; O’Regan & Kitchener, 2005).

Looking in detail at specific nutrients, as opposed to broad dietary groups, most previous studies suggest that carbohydrates are the main factor contributing to calculus formation (Buckley et al., 2014; Radini et al., 2017), as they provide the substrate for the salivary amylase that breaks down sugars in the saliva and enables formation of a film on the hard tooth surface, to which bacteria may adhere. In line with this, we found the highest rates of DC formation in species with a diet consisting of >70% fibers, sugars, and starches (Fig. 4, right contour bound). Species with high protein diets (Fig. 4, left contour bound) were much less likely to develop dental calculus.

However, the formation of DC is dependent on more than just dietary starches, and indeed we identified many outliers (see Fig. 4). Our results indicate that non-dietary factors may contribute to DC formation. We detected significant phylogenetic clustering at a deeper taxonomic scales but not at the tree tips. This indicates that while there is an underlying phylogenetic structure to DC formation, a given species with calculus is not more likely than random to be sister to another species that also forms calculus. Phylogenetic signal was also detected using continuous DC values. Following this, in the modeling we chose to include taxonomic species and phylogenetic species to represent differences in the phylogenetic depth of the DC formation. Similar to the NRI versus NTI, phylogenetic species refers to the deeper scale of tree-wide phylogenetic distances (quantified by the variance-covariance matrix derived from the phylogenetic tree), while taxonomic species treats each species as a discrete group, agnostic of the degree of their phylogenetic relatedness. We found that the variance in our model was split approximately evenly between these two levels, indicating that while there is phylogenetic structure to how the traits evolved, closely related species in our survey may differ in calculus formation from their closest relatives, possibly resulting from recent dietary changes. This result is exemplified in e.g. the binturong, polar bear, and spiny bandicoot (Fig. S3, *Arctictis binturong, Ursus maritimus, Echymipera kalubu*), which all show a shift in both diet and calculus formation compared to their nearest relative.

Other phylogenetically structured morphological factors may play a role in DC formation. Both the NRI and our modeling show phylogenetic signal at a deeper level, such as that between orders, partially explaining patterns in DC formation. The biological and ecological factors that contribute to this are likely as varied as mammalian oral physiology itself: salivary gland activity and position range widely in mammals and salivary composition is known to contribute to DC formation (Hernández-Castañeda et al., 2015; Lieverse, 1999).

Carnivores and aquatic mammals have the smallest salivary glands; seals have very small salivary glands and no parotid salivary glands (Tucker, 1958). Development of the glands varies as well, with large parotid glands common in herbivores (Hofmann et al., 2008). The parotid gland is the largest and most important salivary gland in ruminants, providing half the salivary flow (Kay, 1987). In most ruminants (including several species we surveyed), the opening of the parotid salivary duct is located opposite the first upper molar (Hofmann et al., 2008; Kay, 1987; Nourinezhad et al., 2021), potentially accounting for the high levels of calculus encrustation on the upper jaws of bovids and other ruminants. Although we did not detect significant differences in DC scores between lower and upper jaw across all species, bovids show higher molar DC score on the upper jaw compared to the lower jaw (Fig. 6A).

**Figure 6:**
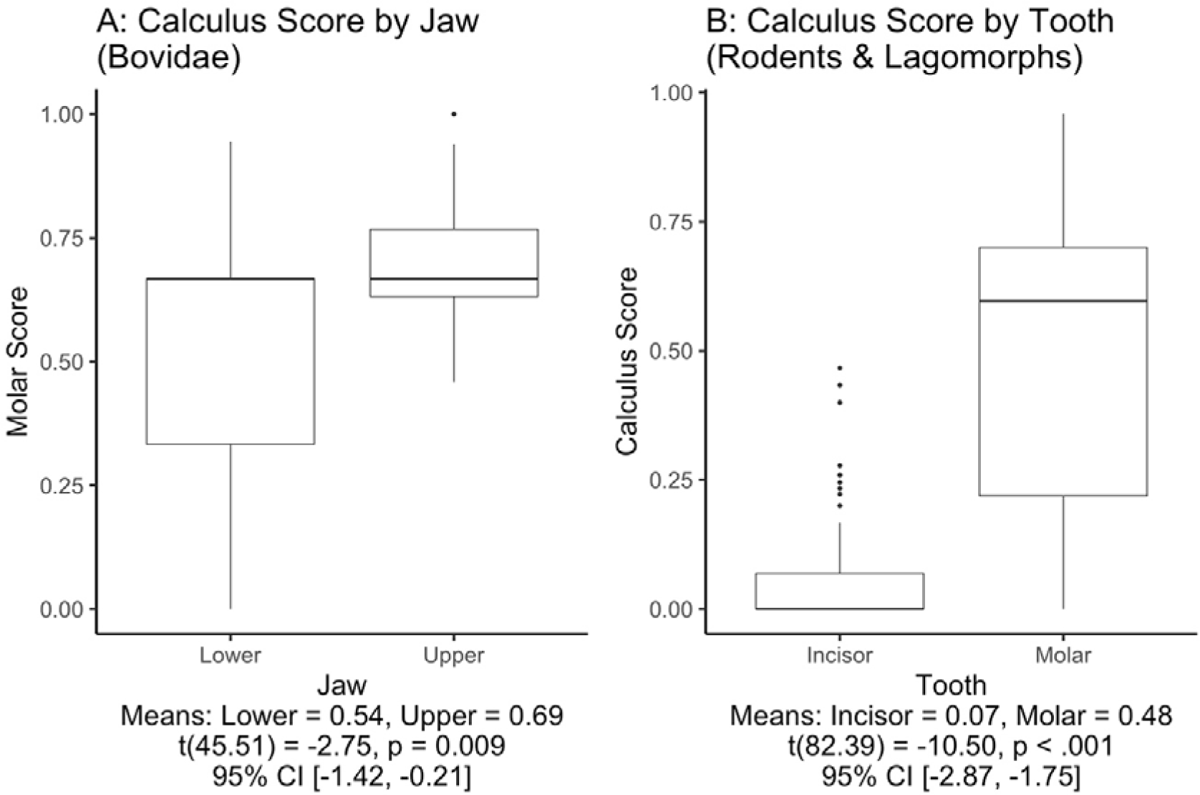
Comparison of calculus scores in bovids and rodents and lagomorphs. Results from Welch’s t-test shown below x axis.

While we focused our analysis primarily on molars because incisors and canines were frequently missing, differences in tooth morphology most certainly affect dental calculus formation. As an example, rodents and lagomorphs have a diet clearly amenable to dental calculus formation. The molars in these mammalian orders consistently showed high levels of encrustation, and the majority of specimens had an average molar score above 0.5 (Fig. 6B). However, calculus was rarely found on incisors; both rodents and lagomorphs possess continuously growing incisors, which likely prevents DC formation through abrasion and loss.

### 4.3 Practical considerations and future perspectives

Shifting priorities and trends in museums’ philosophies of collecting or decisions made by curators long past undoubtedly have an effect on calculus present on skulls today, much as the choices made by curators today will impact research in the years to come. The natural history collections started and amassed largely in Western Europe in the 19^th^ century resulted from a time when scientific priorities lay in exploration and cataloguing. While many questions of taxonomy and systematics the collections were intended for have been resolved, technological developments have repeatedly opened up new avenues of inquiry and expanded the possibilities of what can be done with museum collections (Bein et al., 2025; Brealey et al., 2021; Hahn et al., 2020, 2024; Malakasi et al., 2019; Nakahama, 2021; Nolen et al., 2024; van der Valk et al., 2018; Winker, 2004; Yeates et al., 2016). Long term studies on ecological, evolutionary, and environmental change are especially critical in a time of climate change and biodiversity loss, and historical collections can provide a valuable glimpse into the past, preserving what may be under threat today. Collections, with forward-looking perspectives such as the continued acquisition of contemporary specimens, or a holistic understanding of what constitutes a specimen, inclusive of its parasites, microbiota, environment, etc. (Lendemer et al., 2020; Spellman, 2019) will one day serve future research in ways perhaps unanticipated today (see e.g., “The Next Generation of Natural History Collections”, Schindel & Cook, 2018). Specimen preparation practices can also be rethought to minimize cleaning and maximize preservation of DC, or keeping future sampling in mind when acquiring new specimens, such as prioritizing easy-to-sample lower jaws in ungulates.

Our results show that DC is found in a large diversity of mammals, is primarily present on molars, and generally does not differ in abundance between jaws or tooth sides. Because ease and efficiency of sampling differ between the upper and lower jaws and inner and outer tooth surfaces, these aspects, along with curators’ interests, must be taken into consideration when designing future studies. For most larger species, particularly ungulates, lower jaws are easily handled, frequently collected in large-scale, long-term studies, and can provide a sufficient amount of calculus from 1-2 teeth (Moraitou et al., 2025). Smaller species, while potentially developing large amounts of dental calculus, may not be as easy to sample due to the small size of the skull and fragility of teeth and bones. In addition to historical collections, archeozoological collections can also be sources of DC from much more distant times. Teeth can survive many taphonomic processes and are often found in abundance in excavations, presenting calculus deposits suitable for sampling (see DC on iron age cattle, supplementary Fig. S4).

Using general rules of DC formation that are based on host taxonomy and dietary composition, as derived from our study dataset, we generated predictions across a large number of mammalian species. Our aim is to facilitate research planning for future researchers and projects, and to encourage and inform the use of dental calculus from museum collections. Our predictions show that most mammalian families have an average predicted DC score >0.25. We found species with a score >0.5 to be well suitable for sampling and roughly 0.8 mg of DC material to be sufficient for metagenomic sequencing and analyses (Moraitou et al., 2025). Species with average scores between 0.2 and 0.5 could still yield a suitable amount of calculus, but with lower amounts of encrustation, tooth size and the number of specimens available had a greater influence on the possibility of collecting enough material. Even species with high levels of encrustation may not be suitable for sampling due to their small size (which may require pooling of multiple individuals) or distinctive skull/oral morphology. Our DC predictions can help inform project planning for researchers interested in (preserved) oral microbiomes, by giving an indication of species that most reliably produce dental calculus. Museum curators can also benefit from anticipating which parts of the collection may be of interest to researchers, and help inform collection and acquisition priorities with the future value of DC in mind.

## Acknowledgments

We thank the curators, collection managers, scientific assistants, and others whose assistance was invaluable: at the Naturhistorisches Museum Wien: Frank Zachos, curator of the mammalian collection; Alex Bibl, collection manager; Tim Langnitschke, scientific assistant. At the Africa Museum, Tervuren: Dominique Fonck and Mathys Rotonda. At the Royal Belgian Institute of Natural Sciences: Sébastien Bruaux and Virginie Grignet. We thank Megan Grant from the University of Edinburgh (United Kingdom) for helping gather mammal dietary data, Chanah Bolyos for assistance in specimen scoring and photography, and Ally Phillimore for helpful discussions about statistical analyses. This work was supported by the Swedish Research Council (Formas) grant 2019-00275 to KG and a Darwin Trust studentship for JR.

## Supplementary Figures

**Figure S1:**
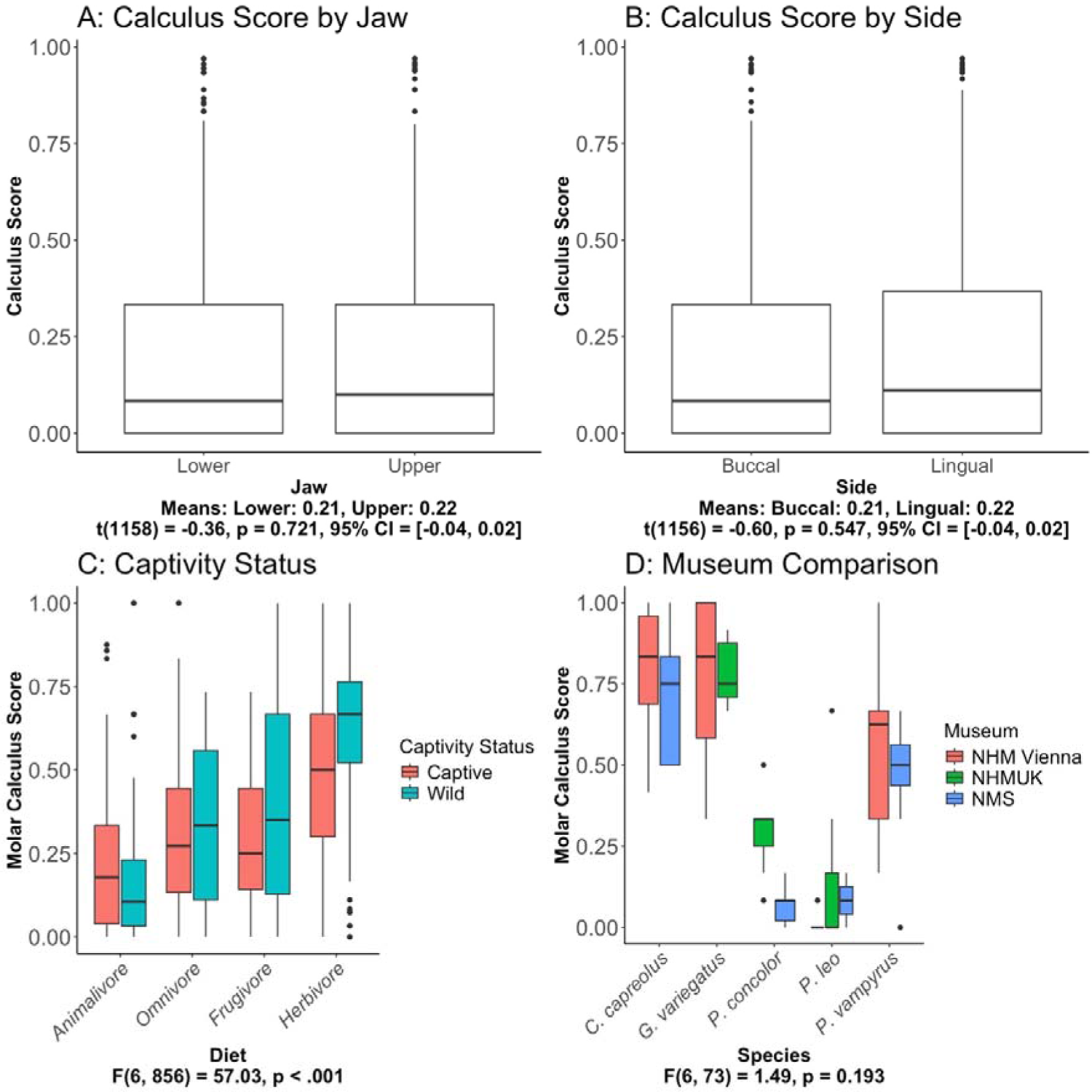
Comparison of calculus scores by jaw (A), side (B), captivity status (C), and museum (D). Y axes show values for DC scores, considering only molars in C and D. Results for ANOVAs (one-way: A & B, nested: C & D) shown below x axis labels. In D, considered species are *Capreolus capreolus* (roe deer, herbivore), *Galeopterus variegatus* (Sunda colugo, herbivore), *Panthera leo* (lion, animalivore), *Pteropus vampyrus* (large flying fox, frugivore) and *Puma concolor* (puma, animalivore).

**Figure S2:**
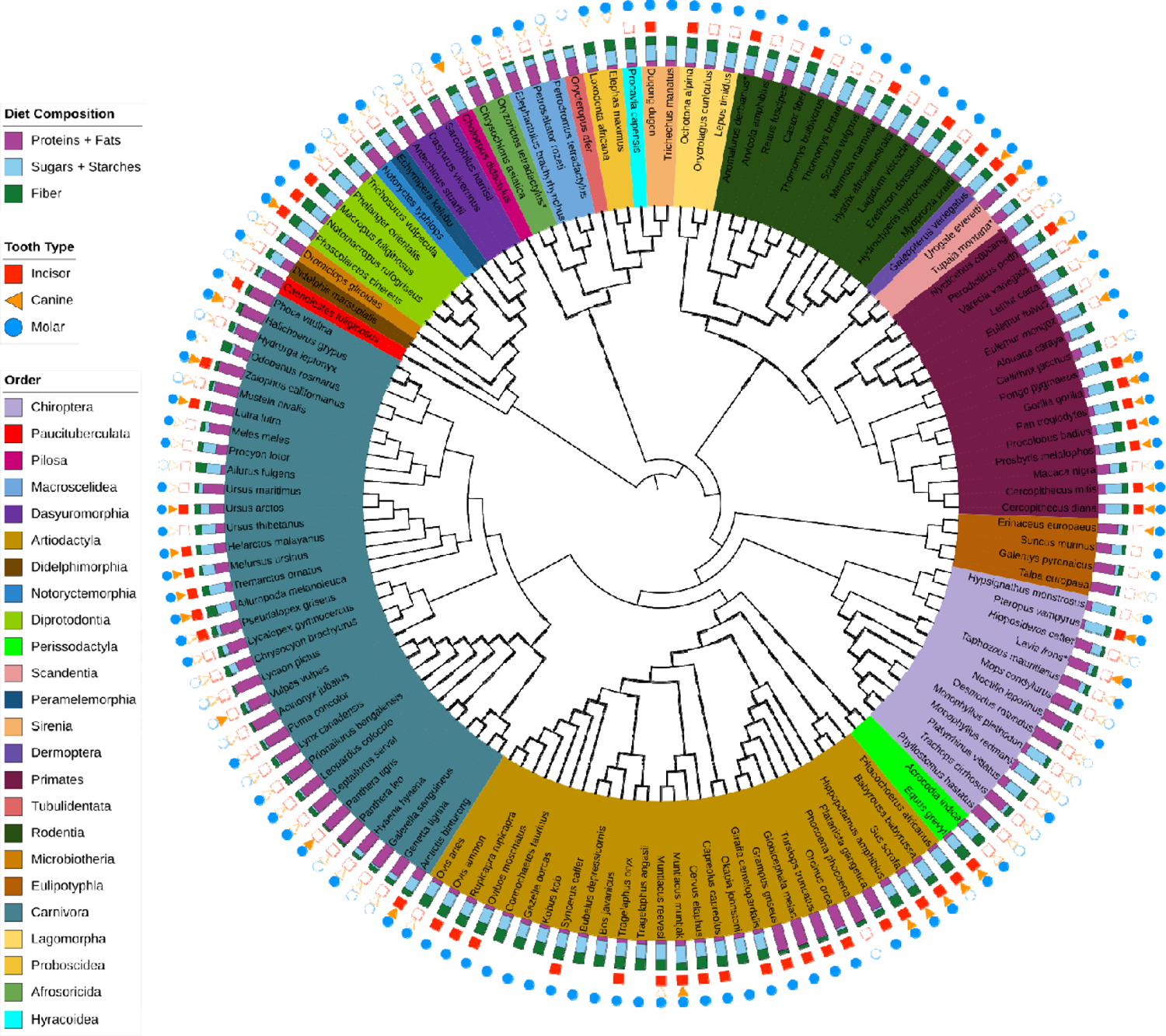
Dental calculus presence across the mammalian phylogeny and tooth types, showing all surveyed wild species (n=141). Mean scores per tooth type were binarized, with a score >0.1 considered as DC presence and shown with a filled shapes along the three outer rings. Unfilled shapes indicate a mean score ≤0.1, considered as absence of DC. No shape indicates that the tooth type was absent for this species. All cetacean teeth were scored as molars. This tree was downloaded from vertlife.org (Upham et al., 2019) using the MCC consensus tree of DNA-only records. Asterisks indicate surveyed species not present in the downloaded tree. For *Anomalurus derbianus, Tupaia montana,* and *Oryzorictes tetradactylus* we used congeneric species as representatives: *A. beecrofti, T. glis, O. hova*.

**Figure S3:**
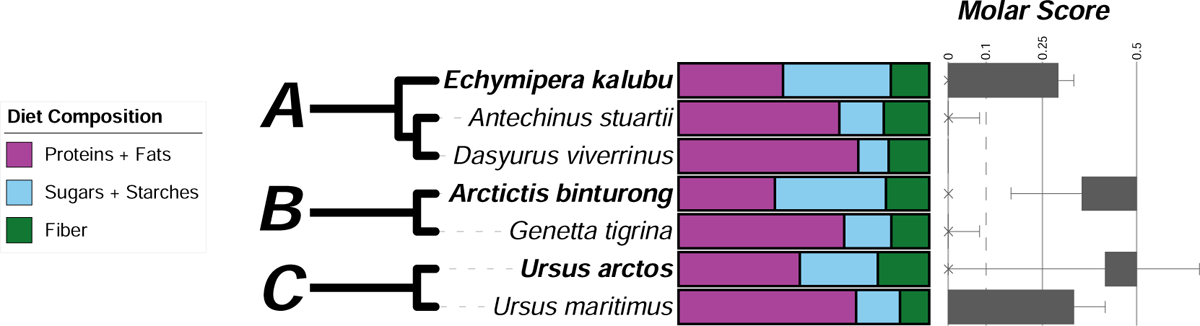
Differences in calculus scores accompanying diet changes in close relatives. Examples taken from the binarized survey phylogeny in Fig. 2. Note the changes in diet (specifically reduced protein intake [purple] and increased fibers, sugars, and starches) relative to their closest surveyed relatives in spiny bandicoot *Echymipera kalubu* (A), binturong *Arctictis binturong* (B), and brown bear *Ursus arctos* (C) and accompanying increase in DC formation.

**Figure S4:**
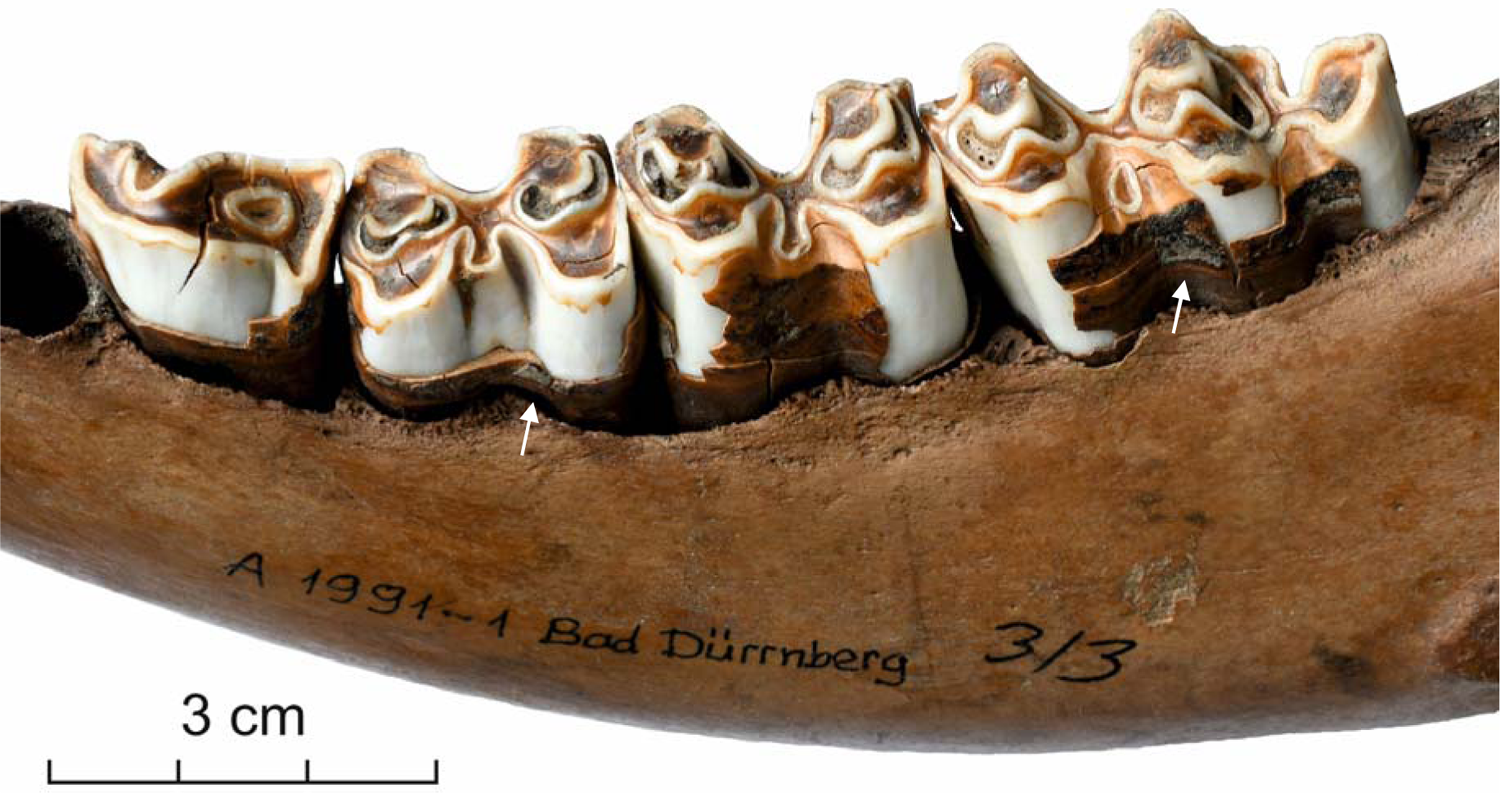
Dental calculus found on Iron Age cattle jaw. Specimen recovered from Bad Dürrnberg, Austria. Calculus deposits indicated with white arrows. Note that the smooth brown layer on the teeth corresponds to enamel and not calculus, with calculus showing rough, whitish surface. Specimen is dated to the Early La Tène to Late La Tène period (c. 450 B.C. to 1 B.C.). Specimen taken from the Archeozoological collection of NHM Vienna (Inventory number: A 1991-1).

## References

Adler, C. J., Dobney, K., Weyrich, L. S., Kaidonis, J., Walker, A. W., Haak, W., A Bradshaw, C. J., Townsend, G., Sołtysiak, A., Alt, K. W., Parkhill, J., & Cooper, A. (2013). Sequencing ancient calcified dental plaque shows changes in oral microbiota with dietary shifts of the Neolithic and Industrial revolutions. Nature Publishing Group. 10.1038/ng.2536

Akcalı, A., & Lang, N. P. (2018). Dental calculus: The calcified biofilm and its role in disease development. Periodontology 2000, 76(1), 109–115. 10.1111/prd.12151

Armitage, P. L. (1975). The extraction and identification of opal phytoliths from the teeth of ungulates. Journal of Archaeological Science, 2(3), 187–197. 10.1016/0305-4403(75)90056-4

Bein, B., Chrysostomakis, I., Arantes, L. S., Brown, T., Gerheim, C., Schell, T., Schneider, C., Leushkin, E., Chen, Z., Sigwart, J., Gonzalez, V., Wong, N. L. W. S., Santos, F. R., Blom, M. P. K., Mayer, F., Mazzoni, C. J., Böhne, A., Winkler, S., Greve, C., & Hiller, M. (2025). Long-read sequencing and genome assembly of natural history collection samples and challenging specimens. Genome Biology, 26(1), 25. 10.1186/s13059-025-03487-9

Brealey, J. C., Leitão, H. G., Hofstede, T., Kalthoff, D. C., & Guschanski, K. (2021). The oral microbiota of wild bears in Sweden reflects the history of antibiotic use by humans. Current Biology, 31(20), 4650–4658.e6. 10.1016/j.cub.2021.08.010

Brealey, J. C., Leitão, H. G., van der Valk, T., Xu, W., Bougiouri, K., Dalén, L., & Guschanski, K. (2020). Dental Calculus as a Tool to Study the Evolution of the Mammalian Oral Microbiome. Molecular Biology and Evolution, 37(10), 3003–3022. 10.1093/molbev/msaa135

Buckley, S., Usai, D., Jakob, T., Radini, A., & Hardy, K. (2014). Dental Calculus Reveals Unique Insights into Food Items, Cooking and Plant Processing in Prehistoric Central Sudan. PLOS ONE, 9(7), e100808. 10.1371/journal.pone.0100808

Ciochon, R. L., Piperno, D. R., & Thompson, R. G. (1990). Opal phytoliths found on the teeth of the extinct ape *Gigantopithecus blacki*: Implications for paleodietary studies. Proceedings of the National Academy of Sciences, 87(20), 8120–8124. 10.1073/pnas.87.20.8120

Clarke, D., & Cameron, A. (1998). Relationship between diet, dental calculus and periodontal disease in domestic and feral cats in Australia. Australian Veterinary Journal, 76(10), 690–693. 10.1111/j.1751-0813.1998.tb12284.x

Coppi, A., Lazzaro, L., Ampoorter, E., Baeten, L., Verheyen, K., & Selvi, F. (2019). Understorey phylogenetic diversity in thermophilous deciduous forests: Overstorey species identity can matter more than species richness. Forest Ecosystems, 6(1), 37. 10.1186/s40663-019-0191-1

Cowan, P. D., Hel-, M. R., Morlon, H., Webb, O., & Kembel, M. S. W. (2020). Package ‘ picante.’ 10.1093/bioinformatics/btq166>.License

Crates, R., Stojanovic, D., & Heinsohn, R. (2023). The phenotypic costs of captivity. Biological Reviews, 98(2), 434–449. 10.1111/brv.12913

Culver, M. (2000). Genomic ancestry of the American puma (*Puma concolor*). Journal of Heredity, 91(3), 186–197. 10.1093/jhered/91.3.186

De La Fuente, C., Flores, S., & Moraga, M. (2013). Dna from human ancient bacteria: A novel source of genetic evidence from archaeological dental calculus. Archaeometry, 55(4), 767–778. 10.1111/j.1475-4754.2012.00707.x

Dzierga, B. M. (2014). Zutrition: Analyzing and Evaluating Diets Fed to Captive Mammals at Capron Park Zoo.

Evans, A. R., & Pineda-Munoz, S. (2018). Inferring Mammal Dietary Ecology from Dental Morphology. In D. A. Croft, D. F. Su, & S. W. Simpson (Eds.), Methods in Paleoecology: Reconstructing Cenozoic Terrestrial Environments and Ecological Communities (pp. 37–51). Springer International Publishing. 10.1007/978-3-319-94265-0_4

Fellows Yates, J. A., Velsko, I. M., Aron, F., Posth, C., Hofman, C. A., Austin, R. M., Parker, C. E., Mann, A. E., Nägele, K., Arthur, K. W., Arthur, J. W., Bauer, C. C., Crevecoeur, I., Cupillard, C., Curtis, M. C., Dalén, L., Díaz-Zorita Bonilla, M., Díez Fernández-Lomana, J. C., Drucker, D. G., … Warinner, C. (2021). The evolution and changing ecology of the African hominid oral microbiome. Proceedings of the National Academy of Sciences, 118(20), e2021655118. 10.1073/pnas.2021655118

Fens, A., & Marcus Clauss, M. C. (2024). Nutrition as an integral part of behavioural management of zoo animals. Journal of Zoo and Aquarium Research, 12(4), 196–204. 10.19227/jzar.v12i4.786

Gittleman, J. L., & Kot, M. (1990). Adaptation: Statistics and a Null Model for Estimating Phylogenetic Effects. Systematic Biology, 39(3), 227–241. 10.2307/2992183

Hahn, E. E., Grealy, A., Alexander, M., & Holleley, C. E. (2020). Museum Epigenomics: Charting the Future by Unlocking the Past. Trends in Ecology & Evolution, 35(4), 295–300. 10.1016/j.tree.2019.12.005

Hahn, E. E., Stiller, J., Alexander, M. R., Grealy, A., Taylor, J. M., Jackson, N., Frere, C. H., & Holleley, C. E. (2024). Century-old chromatin architecture revealed in formalin-fixed vertebrates. Nature Communications, 15(1), 6378. 10.1038/s41467-024-50668-4

Hendy, J., Warinner, C., Bouwman, A., Collins, M. J., Fiddyment, S., Fischer, R., Hagan, R., Hofman, C. A., Holst, M., Chaves, E., Klaus, L., Larson, G., Mackie, M., McGrath, K., Mundorff, A. Z., Radini, A., Rao, H., Trachsel, C., Velsko, I. M., & Speller, C. F. (2018). Proteomic evidence of dietary sources in ancient dental calculus. Proceedings of the Royal Society B: Biological Sciences, 285(1883), 20180977. 10.1098/rspb.2018.0977

Hernández-Castañeda, A. A., Aranzazu-Moya, G. C., Mora, G. M., & de Queluz, D. P. (2015). Chemical salivary composition and its relationship with periodontal disease and dental calculus. Brazilian Journal of Oral Sciences, 14, 159–165. 10.1590/1677-3225v14n2a12

Hofmann, R. R., Streich, W. J., Fickel, J., Hummel, J., & Clauss, M. (2008). Convergent evolution in feeding types: Salivary gland mass differences in wild ruminant species. Journal of Morphology, 269(2), 240–257. 10.1002/jmor.10580

Jin, Y., & Yip, H.-K. (2002). Supragingival calculus: Formation and control. Crit. Rev. Oral Biol. Med, 13, 426–441.

Kapoor, V., Antonelli, T., Parkinson, J. A., & Hartstone-Rose, A. (2016). Oral health correlates of captivity. Research in Veterinary Science, 107, 213–219. 10.1016/j.rvsc.2016.06.009

Kay, R. N. B. (1987). Weights of salivary glands in some ruminant animals. Journal of Zoology, 211(3), 431–436. 10.1111/j.1469-7998.1987.tb01544.x

Keck, F., Rimet, F., Bouchez, A., & Franc, A. (2016). Phylosignal: An R package to measure, test, and explore the phylogenetic signal. Ecology and Evolution, 6(9), 2774–2780. 10.1002/ece3.2051

Kerr, K. R., Morris, C. L., Burke, S. L., & Swanson, K. S. (2013). Apparent total tract macronutrient and energy digestibility of 1-to-3-day-old whole chicks, adult ground chicken, and extruded and canned chicken-based diets in African wildcats (*Felis silvestris lybica*). Zoo Biology, 32(5), 510–517. 10.1002/zoo.21084

Lendemer, J., Thiers, B., Monfils, A. K., Zaspel, J., Ellwood, E. R., Bentley, A., LeVan, K., Bates, J., Jennings, D., Contreras, D., Lagomarsino, L., Mabee, P., Ford, L. S., Guralnick, R., Gropp, R. E., Revelez, M., Cobb, N., Seltmann, K., & Aime, M. C. (2020). The Extended Specimen Network: A Strategy to Enhance US Biodiversity Collections, Promote Research and Education. BioScience, 70(1), 23–30. 10.1093/biosci/biz140

Lieverse, A. R. (1999). Diet and the aetiology of dental calculus. International Journal of Osteoarchaeology, 9(4), 219–232. 10.1002/(SICI)1099-1212(199907/08)9:4<219::AID-OA475>3.0.CO;2-V

Lintulaakso, K., Tatti, N., & Žliobaitė, I. (2023). Quantifying mammalian diets. Mammalian Biology, 103(1), 53–67. 10.1007/s42991-022-00323-6

Mackie, M., Hendy, J., Lowe, A. D., Sperduti, A., Holst, M., Collins, M. J., & Speller, C. F. (2017). Preservation of the metaproteome: Variability of protein preservation in ancient dental calculus. STAR: Science & Technology of Archaeological Research, 3(1), 58–70. 10.1080/20548923.2017.1361629

Malakasi, P., Bellot, S., Dee, R., & Grace, O. M. (2019). Museomics Clarifies the Classification of *Aloidendron* (Asphodelaceae), the Iconic African Tree Aloes. Frontiers in Plant Science, 10(October), 1–11. 10.3389/fpls.2019.01227

Mann, A. E., Sabin, S., Ziesemer, K., Vågene, Å. J., Schroeder, H., Ozga, A. T., Sankaranarayanan, K., Hofman, C. A., Fellows Yates, J. A., Salazar-García, D. C., Frohlich, B., Aldenderfer, M., Hoogland, M., Read, C., Milner, G. R., Stone, A. C., Lewis, C. M., Krause, J., Hofman, C., … Warinner, C. (2018). Differential preservation of endogenous human and microbial DNA in dental calculus and dentin. Scientific Reports, 8(1), 9822. 10.1038/s41598-018-28091-9

Moraitou, M., Forsythe, A., Fellows Yates, J. A., Brealey, J. C., Warinner, C., & Guschanski, K. (2022). Ecology, not host phylogeny, shapes the oral microbiome in closely related species. *Molecular Biology and Evolution*, msac263. 10.1093/molbev/msac263

Moraitou, M., Richards, J., Bolyos, C., Saliari, K., Gilissen, E., Timmons, Z., Kitchener, A. C., Pauwels, O. S. G., Sabin, R., Kokkini, P., Miguez, R. P., & Guschanski, K. (2025). Host traits impact the outcome of metagenomic library preparation from dental calculus samples across diverse mammals (p. 2025.03.19.643754). bioRxiv. 10.1101/2025.03.19.643754

Nakahama, N. (2021). Museum specimens: An overlooked and valuable material for conservation genetics. Ecological Research, 36(1), 13–23. 10.1111/1440-1703.12181

Nolen, Z. J., Jamelska, P., Lara, A. S. T., Wahlberg, N., & Runemark, A. (2024). Species-specific loss of genetic diversity and exposure of deleterious mutations following agricultural intensification. bioRxiv, 2024–10.

Nourinezhad, J., Moarabi, A., & Ahkalani, M. S. R. (2021). Detailed gross anatomic and sialographic characteristics of major salivary glands in water buffaloes (*Bubalus bubalis*). Anatomical Science International, 96(3), 427–442. 10.1007/s12565-021-00609-8

O’Regan, H. J., & Kitchener, A. C. (2005). The effects of captivity on the morphology of captive, domesticated and feral mammals. Mammal Review, 35(3/4), 215–230. 10.1111/j.1365-2907.2005.00070.x

Ozga, A. T., & Ottoni, C. (2023). Dental calculus as a proxy for animal microbiomes. Quaternary International, *653–654*, 47–52. 10.1016/j.quaint.2021.06.012

Pagel, M. (1999). Inferring the historical patterns of biological evolution. Nature, 401(October 1999), 877–884.

Power, R. C., Salazar-García, D. C., Straus, L. G., González Morales, M. R., & Henry, A. G. (2015). Microremains from El Mirón Cave human dental calculus suggest a mixed plant– animal subsistence economy during the Magdalenian in Northern Iberia. Journal of Archaeological Science, 60, 39–46. 10.1016/j.jas.2015.04.003

Power, R. C., Salazar-García, D. C., Wittig, R. M., Freiberg, M., & Henry, A. G. (2015). Dental calculus evidence of Taï Forest Chimpanzee plant consumption and life history transitions. Scientific Reports, 5(1), 15161. 10.1038/srep15161

Putrino, A., Marinelli, E., Galeotti, A., Ferrazzano, G. F., Ciribè, M., & Zaami, S. (2024). A Journey into the Evolution of Human Host-Oral Microbiome Relationship through Ancient Dental Calculus: A Scoping Review. Microorganisms, 12(5), Article 5. 10.3390/microorganisms12050902

Radini, A., Nikita, E., Buckley, S., Copeland, L., & Hardy, K. (2017). Beyond food: The multiple pathways for inclusion of materials into ancient dental calculus. American Journal of Physical Anthropology, 162(S63), 71–83. 10.1002/ajpa.23147

Reuter, D. M., Hopkins, S. S. B., & Price, S. A. (2023). What is a mammalian omnivore? Insights into terrestrial mammalian diet diversity, body mass and evolution. Proceedings of the Royal Society B: Biological Sciences, 290(1992), 20221062. 10.1098/rspb.2022.1062

Rivera-Perez, J. I., Cano, R. J., Narganes-Storde, Y., Chanlatte-Baik, L., & Toranzos, G. A. (2015). Retroviral DNA Sequences as a Means for Determining Ancient Diets. PLOS ONE, 10(12), e0144951. 10.1371/journal.pone.0144951

Schindel, D. E., & Cook, J. A. (2018). The next generation of natural history collections. PLOS Biology, 16(7), e2006125. 10.1371/journal.pbio.2006125

Shaiber, A., Willis, A. D., Delmont, T. O., Roux, S., Chen, L.-X., Schmid, A. C., Yousef, M., Watson, A. R., Lolans, K., Esen, Ö. C., Lee, S. T. M., Downey, N., Morrison, H. G., Dewhirst, F. E., Mark Welch, J. L., & Eren, A. M. (2020). Functional and genetic markers of niche partitioning among enigmatic members of the human oral microbiome. Genome Biology, 21(1), 292. 10.1186/s13059-020-02195-w

Spellman, G. M. (2019). The Extended Specimen: Emerging Frontiers in Collections-Based Ornithological Research. The Auk, 136(3), ukz024. 10.1093/auk/ukz024

Tucker, R. (1958). Taxonomy of the Salivary Glands of Vertebrates. Systematic Zoology, 7(2), 74–83. 10.2307/2411794

Upham, N. S., Esselstyn, J. A., & Jetz, W. (2019). Inferring the mammal tree: Species-level sets of phylogenies for questions in ecology, evolution, and conservation. PLOS Biology, 17(12), e3000494. 10.1371/journal.pbio.3000494

van der Valk, T., Sandoval-Castellanos, E., Caillaud, D., Ngobobo, U., Binyinyi, E., Nishuli, R., Stoinski, T., Gilissen, E., Sonet, G., Semal, P., Kalthoff, D. C., Dalén, L., & Guschanski, K. (2018). Significant loss of mitochondrial diversity within the last century due to extinction of peripheral populations in eastern gorillas. Scientific Reports, 8(1), 6551. 10.1038/s41598-018-24497-7

Verstraete, F. J. M. (2003). Advances in diagnosis and treatment of small exotic mammal dental disease. Seminars in Avian and Exotic Pet Medicine, 12(1), 37–48. 10.1053/saep.2003.127877

Viechtbauer, W. (2010). Conducting Meta-Analyses in R with the metafor Package. Journal of Statistical Software, 36(3), 1–48. 10.18637/jss.v036.i03

Warinner, C., Rodrigues, J. F. M., Vyas, R., Trachsel, C., Shved, N., Grossmann, J., Radini, A., Hancock, Y., Tito, R. Y., & Fiddyment, S. (2014). Pathogens and host immunity in the ancient human oral cavity. Nat. Genet, 46, 336–344.

Warinner, C., Speller, C., & Collins, M. J. (2015). A new era in palaeomicrobiology: Prospects for ancient dental calculus as a long-term record of the human oral microbiome. Philosophical Transactions of the Royal Society B: Biological Sciences, 370(1660). 10.1098/rstb.2013.0376

Whitehouse-Tedd, K. M., Lefebvre, S. L., & Janssens, G. P. J. (2015). Dietary Factors Associated with Faecal Consistency and Other Indicators of Gastrointestinal Health in the Captive Cheetah (*Acinonyx jubatus*). PLOS ONE, 10(4), e0120903. 10.1371/journal.pone.0120903

Winker, K. (2004). Natural History Museums in a Postbiodiversity Era. BioScience, 54(5), 455–459. 10.1641/0006-3568(2004)054[0455:NHMIAP]2.0.CO;2

Yaprak, E., Yolcubal, İ., Sinanoğlu, A., Doğrul-Demiray, A., Guzeldemir-Akcakanat, E., & Marakoğlu, İ. (2017). High levels of heavy metal accumulation in dental calculus of smokers: A pilot inductively coupled plasma mass spectrometry study. Journal of Periodontal Research, 52(1), 83–88. 10.1111/jre.12371

Yeates, D. K., Zwick, A., & Mikheyev, A. S. (2016). Museums are biobanks: Unlocking the genetic potential of the three billion specimens in the world’s biological collections. Current Opinion in Insect Science, 18, 83–88. 10.1016/j.cois.2016.09.009

